# Sodium valproate rescues expression of *TRANK1* in iPSC-derived neural cells that carry a genetic variant associated with serious mental illness

**DOI:** 10.1101/192815

**Authors:** Xueying Jiang, Sevilla D Detera-Wadleigh, Nirmala Akula, Barbara S. Mallon, Liping Hou, Tiaojiang Xiao, Gary Felsenfeld, Xinglong Gu, Francis J. McMahon

## Abstract

Biological characterization of genetic variants identified in genome-wide association studies (GWAS) remains a substantial challenge. Here we used human induced pluripotent stem cells (hiPSC) and their neural derivatives to characterize common variants on chromosome 3p22 that have been associated by GWAS with major mental illnesses. HiPSC-derived neural progenitor cells carrying the risk allele of the single nucleotide polymorphism (SNP), rs9834970, displayed lower baseline *TRANK1* expression that was rescued by chronic treatment with therapeutic dosages of valproic acid (VPA). Although rs9834970 has no known function, we demonstrated that a nearby SNP, rs90832, strongly affects binding by the transcription factor, CTCF, and that the high-affinity allele usually occurs on haplotypes carrying the rs9834970 risk allele. Decreased expression of *TRANK1* perturbed expression of many genes involved in neural development and differentiation. These findings have important implications for the pathophysiology of major mental illnesses and the development of novel therapeutics.

**Highlights:**

- hiPSC-derived neural cells carrying a mental illness risk allele showed lower expression of *TRANK1*
- Valproic acid rescued *TRANK1*expression in cells carrying the risk allele
- Risk haplotypes usually carry an allele that increased CTCF binding
- Reduced expression of *TRANK1* perturbed genes involved in neural development and differentiation

**In Brief:** Using neural derivatives of human induced pluripotent stem cells, Jiang et al. demonstrate that genetic variants associated with mental illness alter transcription factor binding and decrease expression of a nearby gene, an effect which is rescued by valproic acid.

## INTRODUCTION

Genome-wide association studies (GWASs) of bipolar disorder (BD) and schizophrenia have identified several reproducible genetic markers, but the biological consequences of most variants remain undefined (Richards et al., 2012; Shinozaki and Potash, 2014). Many common complex traits are mediated by expression quantitative trait loci (eQTLs) that affect expression of nearby (cis) or distant (trans) genes (Kirsten et al., 2015). Altered gene expression is thus one of the most important mechanisms underlying such genotype-phenotype associations (GTex Consortium, 2013). However, eQTLs may act only in specific tissues (De Gobbi et al., 2007; Liu et al., 2014; Mele et al., 2015), or during particular stages of development (Gay et al., 2015; Latham, 1995). Such tissue and time-specific effects complicate studies that seek to assess the impact of allelic variation on gene expression in the human central nervous system (CNS), where appropriate cells from living individuals are difficult or impossible to obtain. Past eQTL studies of variants implicated in brain disorders have thus depended on limited supplies of human postmortem brain. Such studies are valuable, but cannot directly assess developmental effects and are often confounded by age, gender, treatment history, drug or alcohol abuse, and agonal events (Rueckert et al., 2013).

Induced pluripotent stem cells (iPSC) offer an attractive alternative approach (Korecka et al., 2016). eQTL studies in human iPSC-derived cells exploit a renewable supply of cells largely free from the complications of postmortem tissue. iPSC can be easily generated from patients of known genetic background, further reducing sources of variance that can obscure subtle signals. Model systems based on iPSC-derived cells can follow changes in gene expression during development and can also be experimentally manipulated with drugs, toxins, or gene-editing techniques. While iPSC-derived cells can be affected by somatic changes in DNA sequence or copy number (McConnell et al., 2013) and do not necessarily reflect epigenetic marks in the donor’s brain (Kim et al., 2010), they nevertheless offer great potential for dissecting the molecular mechanisms of complex brain disorders.

In this study, we used iPSC and their neural derivatives to investigate the cellular impact of common genetic polymorphisms previously associated with bipolar disorder (BD) and schizophrenia by genome-wide association studies (GWAS). The polymorphisms lie on chromosome 3p22 near the gene *TRANK1* (also known as *LBA1*). *TRANK1* encodes tetratricopeptide repeat and ankyrin repeat containing 1, a protein that is expressed in the brain and several other tissues, but whose function remains unknown. A common variant (rs9834970) located some 15 kb 3’ of *TRANK1* has shown genome-wide significant association with BD in several GWAS (Chen et al., 2013a; Goes et al., 2012; Ruderfer et al., 2013), and nearby markers have been associated with schizophrenia and other mental illnesses (Cross-Disorder Group of the Psychiatric Genomics Consortium, 2013; Schizophrenia Working Group of the Psychiatric Genomics Consortium, 2014). We have previously shown that valproic acid (VPA), an effective treatment for BD, increases *TRANK1* expression in immortalized cell lines (Chen et al., 2013a), but the relationship between VPA exposure, genetic risk variants, and expression of *TRANK1* in neural cells has remained unclear.

The present study had several objectives: 1) to assess the impact of common variants within the 3p22 GWAS locus on *TRANK1* expression in patient-derived neural cells; 2) to assess the effects of treatment with VPA and another commonly-used therapeutic agent, lithium; and 3) to uncover genes and gene networks whose expression correlates with that of *TRANK1*, thereby shedding new light on its function. The findings establish that genetic variants on chromosome 3p22 associated with risk for major mental illnesses alter binding of the transcription factor, CTCF, and decrease expression of *TRANK1* in neural cells. Furthermore, chronic treatment with therapeutic dosages of VPA restored expression of *TRANK1* mRNA and protein to control levels. These findings have important implications for the pathophysiology of major mental illnesses and the development of novel therapeutics.

## RESULTS

### Induction and characterization of human iPSCs and their neural derivatives

iPSC lines were generated from fibroblasts using standard lentiviral transduction procedures (see Methods). All lines exhibited typical stem cell morphology. Immunofluorescence analysis demonstrated expression of the pluripotency markers OCT4 (>99%), Nanog (>97%), Tra-1-81 (>95%), and Tra-1-60 (>99%) (Figure 1A). Subsequent flow cytometry analysis showed absent expression of SSEA1 and increased expression of Tra-1-60, Tra-1-81, OCT4, and SSEA-4 (Figure 1B), consistent with pluripotent status (Yu et al., 2007)).

**Figure 1.**
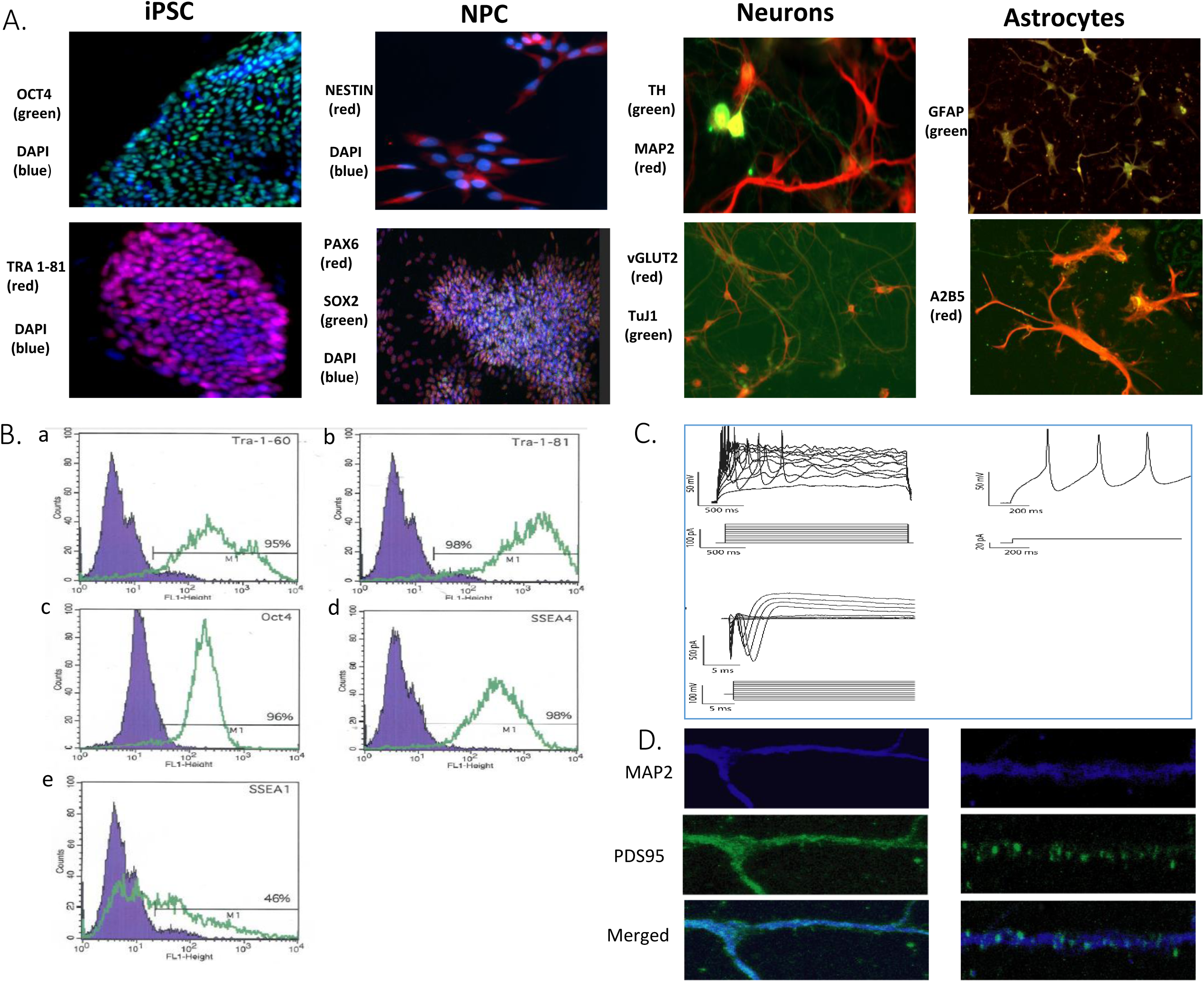
Generation of iPSCs and neural derivatives. (A.) Representative images of induced pluripotent stem cells (iPSCs) and their neural derivatives subjected to immunocytochemical analysis of pluripotency and neural cell differentiation. Far left: iPSCs showed the typical morphology of human embryonic stem cells (hESCs) and positive staining for pluripotency markers OCT4 (green) and DAPI (blue) (upper panel); Tra1-81 (red) and DAPI (blue) (lower panel). Second from left: iPSC-derived neural progenitor cells (NPCs) immunostained positive for neural lineage markers Nestin (red) (upper panel), Pax6 (red) and Sox2 (green) (lower panel). Third from left: iPSC-derived neurons expressed neural markers MAP2 (red), TH (green), (upper panel), β III tubulin (green) and vGlut2 (red) (lower panel). Far right: Astrocytes generated from NPCs exhibited typical astrocyte morphology and expressed astrocyte markers GFAP (green) (upper panel) and A2B5 (red) (lower panel). (B.) Human induced pluripotent stem cells were subjected to flow cytometric to analysis with pluripotency markers. In the merged image, the blue area shows the fluorescence intensity of the IgG negative control antibody, while intensities of the antibodies of interest are shown in green. Results of negative control and iPSC lines are shown in (a) Tra-1 60 (9%, 95%), (b) Tra_1 81 (5%, 98%), (c) OCT4 (4%, 96%), (d) SSEA-4 (3%, 98%), and (e) SSEA-1(10%, 46%). (C.) Representative traces from whole cell-patch clamp recordings in 6-week post-differentiation neurons, demonstrating typical electrophysiological activity. (D.) Immunochemical staining of MAP2 (blue) and PSD95 (green) in 6-week neurons, demonstrating typical synapse staining patterns.

Global gene expression profiles were compared among fibroblasts, iPSCs, iPSC derived neural progenitor cells, and neurons. As expected, fibroblasts, iPSCs, iPSC-derived neural progenitor cells, and neurons formed distinct clusters on the basis of gene expression (Supplemental Figure 1A). All studied iPSC lines exhibited gene expression profiles typical of pluripotent stem cells (Supplemental Figure 1B, S1C). Real-time polymerase chain reaction (RT-PCR) assays demonstrated expression of the pluripotency markers TDGF-1, NANOG, OCT4 and SOX2 (Supplemental Figure 2A) in all studied iPSC lines, increased 100- to 2000-fold relative to the original fibroblasts. G-banding and spectral karyotype analysis demonstrated that all iPSC lines maintained a normal human karyotype (Supplemental Figure 2B).

**Figure 2.**
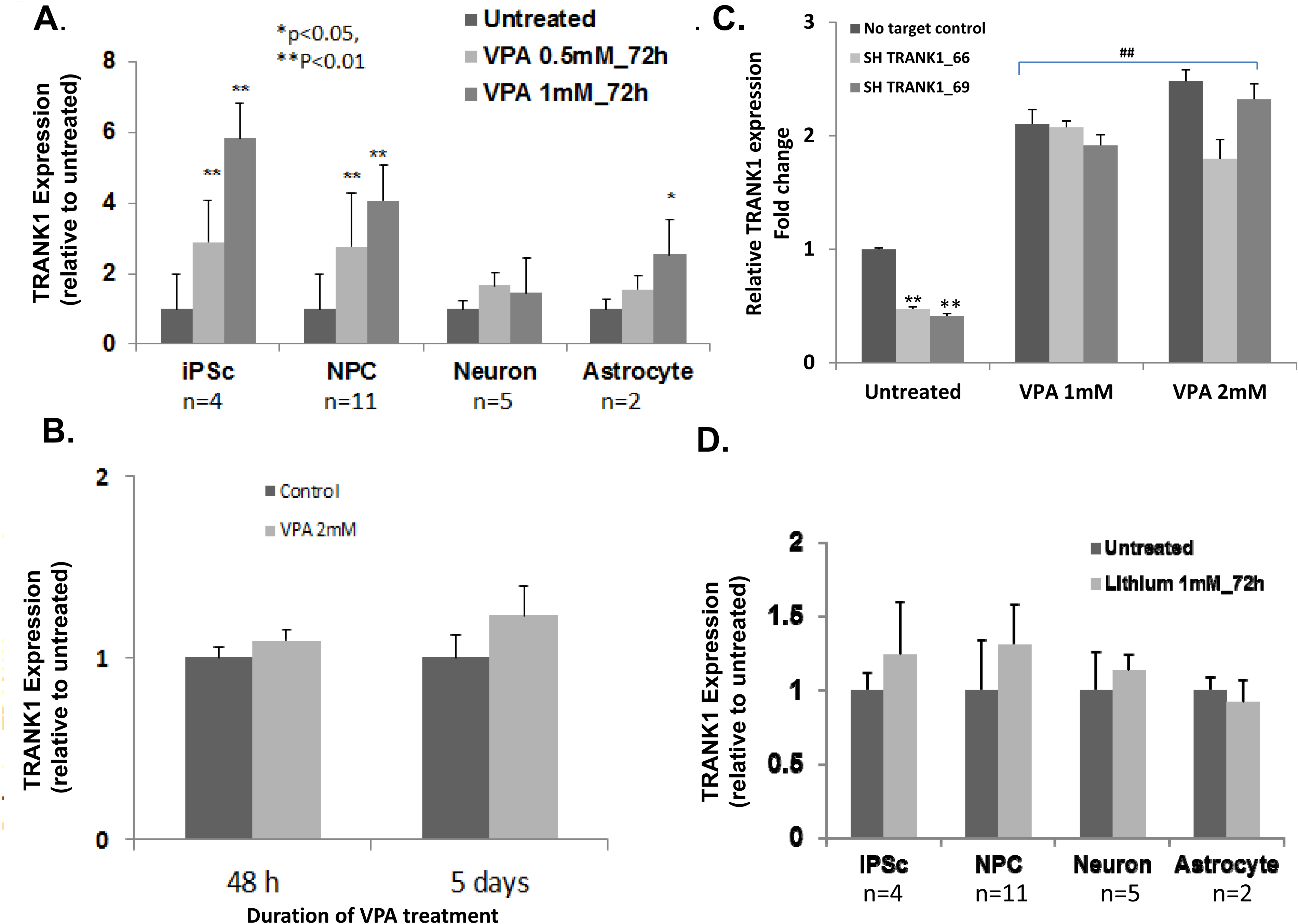
*TRANK1* mRNA expression after VPA or lithium treatment. (A.) Human induced pluripotent stem cell (iPSC), neural progenitor cell (NPC), neuronal, and astrocyte lines were grown in independent cultures and treated with therapeutic dosages of VPA (0.5mM, 1mM) or vehicle for 72 hours. *TRANK1* mRNA expression was determined by reverse transcription polymerase chain reaction(RT-PCR). VPA treatment significantly increased *TRANK1* expression in iPSCs, NPCs, and astrocytes, but not in neurons (*P<0.05, **P <0.01). (B.) *TRANK1* expression in rat E18 hippocampal neurons cultured for 8d before treatment was also unaffected by VPA, even after 48h or 5 days of treatment a 2mM dose. (C.) *TRANK1* expression in HeLa cells was significantly increased by treatment with 1mM or 2mM VPA after stable shRNA knockdown of *TRANK1*. **P<0.01 TRANK1 shRNA knockdown vs no target shRNA control. ## P<0.01 VPA (1mM or 2mM) treated vs untreated condition. D.) Lithium (1mM) had no effect on *TRANK1* expression in any of the 4 cell types tested.

iPSC lines were differentiated into neural progenitor cells (NPCs), neurons, and astrocytes using published protocols (Methods). To confirm a neural lineage, quantitative real time-PCR (qRT–PCR) analysis was carried out for several established NPC marker genes. As expected, the pluripotency-associated genes *NANOG* and *POU5F1* (encoding OCT4) were down-regulated in NPCs, while *PAX6* and *POU3F2*, *CXCR4*, *HES4* were markedly upregulated (Supplemental Figure 2C). All NPC lines were also positive for PAX6, NESTIN and SOX2 immunofluorescence (Figure 1A).

Two NPC lines were differentiated into astrocytes. After seven weeks of differentiation, more than 80% of the cells stained positive for the astrocytic markers glial fibrillary acidic protein (GFAP) and A2B5 (Figure 1A).

Differentiation of five NPC lines into neurons was allowed to proceed for 8 to 12 weeks at which stage cells expressed neuronal antigens, including neuron-specific β-tubulin (TuJ1) and microtubule-associated protein 2 (MAP2). A small proportion of cells stained positive for tyrosine hydroxylase (TH) or vesicular glutamate transporter (VGLUT2), characteristic of a mixed culture of dopaminergic and glutamatergic neurons (Figure 1A). Patch-clamp analysis verified that differentiated neurons displayed membrane potentials, sodium-potassium currents, and action potentials (Figure 1C). These neurons also stained positive for postsynaptic density 95 (PSD95) (Figure 1D). These data are consistent with a functional neuronal phenotype.

### *TRANK1* expression after VPA treatment in iPSCs and their neuronal derivatives

Previously, we showed that therapeutic dosages of the mood stabilizer, VPA, increased expression of *TRANK1* in immortalized, non-neural cell lines (Chen et al 2013a). To assess the effects of VPA on neural cell lines, we treated iPSCs and their neural derivatives with therapeutic dosages of VPA. In both iPSCs and NPCs, *TRANK1* expression increased 3 to 6-fold after 72 hr treatment with VPA (Figure 2A). Expression increased in proportion to VPA dosage over the typical therapeutic range of 0.5 to 1.0 mM. VPA also increased *TRANK1* expression in astrocytes, but the changes, while significant, were less pronounced than those observed in the iPSC and NPC cultures. In contrast, VPA had no detectable effect on *TRANK1* expression in iPSC-derived neurons. A similar lack of change in *TRANK1* expression was seen in cultured rat hippocampal neurons treated for up to 5 days at VPA doses up to 2mM (Figure 2B). These results show that VPA has the greatest effects on *TRANK1* expression at earlier developmental stages, and that these effects diminish with differentiation.

VPA also reversed the effects of two different shTRANK1 knockdown vectors, in HeLa cells (Figure 2C, n=6, **##** p<0.0001 treated vs untreated). This suggests that VPA affects *TRANK1* expression at the level of transcription.

Unlike VPA, but consistent with our published results in immortalized cell lines (Chen et al 2014), the mood stabilizer lithium chloride had no effect on *TRANK1* expression in any of the iPSC-derived cells we tested (Figure 2D). Thus, the subsequent experiments focused exclusively on VPA treatment.

### Genetic variation at rs9834970 and *TRANK1* expression in iPSCs and their neural derivatives

Previous GWAS have consistently found that the G-allele of rs9834970 is significantly more frequent than the A-allele among people with BD or schizophrenia (Bergen et al., 2012; Chen et al., 2013a; Ruderfer et al., 2014a; Shinozaki and Potash, 2014). This SNP lies about 15 kb downstream of *TRANK1*, within a DNaseI hypersensitive site (ENCODE). Conditional analysis of BD GWAS data confirmed that rs9834970 accounted for all of the association signal at this locus (Figure S9). To test whether variation at rs9834970 affects expression of *TRANK1*, we compared *TRANK1* expression in iPSC and NPC lines of known genotype.

Genotype at rs9834870 had a substantial impact on baseline expression of *TRANK1* mRNA in both iPSC and NPC. Cell lines carrying the risk allele (AG or GG genotypes) showed significantly lower baseline *TRANK1* expression than AA homozygotes. In NPCs, carriers of the GG genotype (n=3) showed a 5.6-fold decrease in *TRANK1* expression (p<0.0001) compared to carriers of the AA genotype (n = 3) and a 1.35-fold decrease in *TRANK1* expression (p<0.05) compared to carriers of the AG genotype (n = 5) (Figure 3A, left panel). Similar results were seen in iPSCs, where homozygous risk-allele carriers (GG genotype) showed significantly lower baseline *TRANK1* expression than AA homozygotes (p<0.0001; Figure S3).

**Figure 3.**
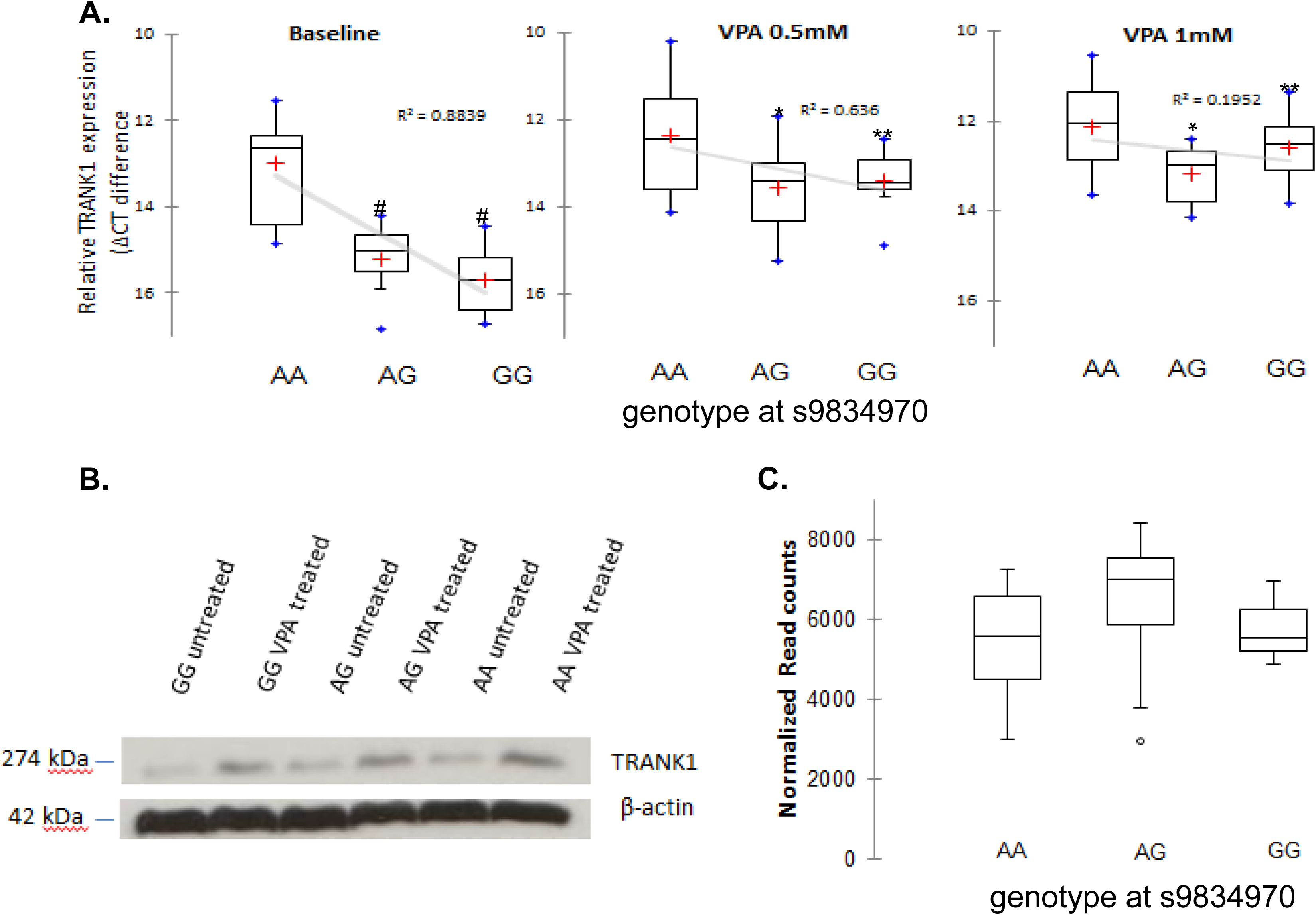
Genotypic effects of rs9834970 on *TRANK1* mRNA and protein expression. **(A.)** Neural progenitor cells (NPCs) carrying the risk allele (G) showed reduced *TRANK1* mRNA expression (left) that was rescued by 72-hr of VPA treatment at dosages of 0.5mM (middle) or 1mM (right). Values are expressed as mean relative δCT difference ± S.E.M. #p<0.0001 baseline AG, GG allele compared to AA allele at rs9834970, *p<0.05, **P<0.01, VPA treatment (0.5mM, 1mM) compared to untreated baseline. Genotypes at rs9834970 and sample sizes: AA = 3, GG=3, AG=5. (B.) Western blot: Qualitative increase in binding of anti-TRANK1 antibody to 274 kDa protein band extracted from NPCs in A-allele carriers and after treatment with 1mM VPA; n =3 for each condition. (C.) No relationship between rs9834970 genotype and *TRANK1* mRNA counts (n=22) in human postmortem dorsolateral prefrontal cortex.

To test whether similar genotype effects could be detected in postmortem brain tissue, we used RNA sequencing data obtained from human postmortem dorsolateral prefrontal cortex (DLPFC) in 22 individuals (Akula et al. 2014). Baseline expression was very low, and no genotype-specific effects on *TRANK1* expression were observed in this brain region (Figure 3C). SNP rs9834970 also showed no evidence of association with *TRANK1* expression in any of the 10 brain regions catalogued in the BRAINEAC database (Kreutzer et al., 2017) (http://www.braineac.org/) (Figure S4), or in any of the neural tissues reported by GTEX (http://www.gtexportal.org, accessed 2/23/2017). These results suggest that rs9834970 either does not affect expression of *TRANK1* in mature brain tissue or exerts cell-type specific effects that cannot be easily detected in homogenated brain tissue.

### VPA Treatment Rescues Expression of *TRANK1* in Risk-Allele Carriers

We next assessed the impact of VPA treatment on *TRANK1* expression. We found that chronic treatment with therapeutic doses of VPA normalized expression of *TRANK1* in NPCs derived from carriers of the risk allele of rs9834970 (8.7 fold increase for GG, p< 0.0001; 3.8 fold increase for AG, p<0.0001, 1.5 fold increase for AA, P<0.05, Figure 3A, middle and right panels). ANOVA analysis confirmed a significant VPA by genotype interaction (F(1,46)=4.452; p<0.05). To test whether the observed changes in mRNA expression corresponded to a change in TRANK1 protein expression, we performed Western blotting in cell lysates derived from NPC lines. This confirmed that TRANK1 protein expression was lower in risk (G) allele carriers than in AA homozygotes, and that VPA treatment increased expression of TRANK1 protein (Fig 3B). We conclude that VPA treatment rescued expression of *TRANK1* mRNA and TRANK1 protein in NPCs that carry the risk allele at rs9834970.

### Predicted regulatory effects of rs9834970 and nearby SNPs

A number of common SNPs near rs9834970 are all associated (in linkage disequilibrium) with one another (Figure 4A). Five of these SNPs have been reported to be associated with BD and/or schizophrenia in previous GWAS (2014; Chen et al., 2013b; Cross-Disorder Group of the Psychiatric Genomics, 2013; Goes et al., 2012; Muhleisen et al., 2014; Ruderfer et al., 2014b; Schizophrenia Working Group of the Psychiatric Genomics, 2014). SNPs in this region form 4 common haplotypes in individuals of European-ancestry from the 1,000 Genomes project (http://www.1000genomes.org/) (Figure 4B). The rs9834970 risk allele is carried by two of these haplotypes (highlighted in yellow), both of which also carry the “G” allele at rs906482.

**Figure 4.**
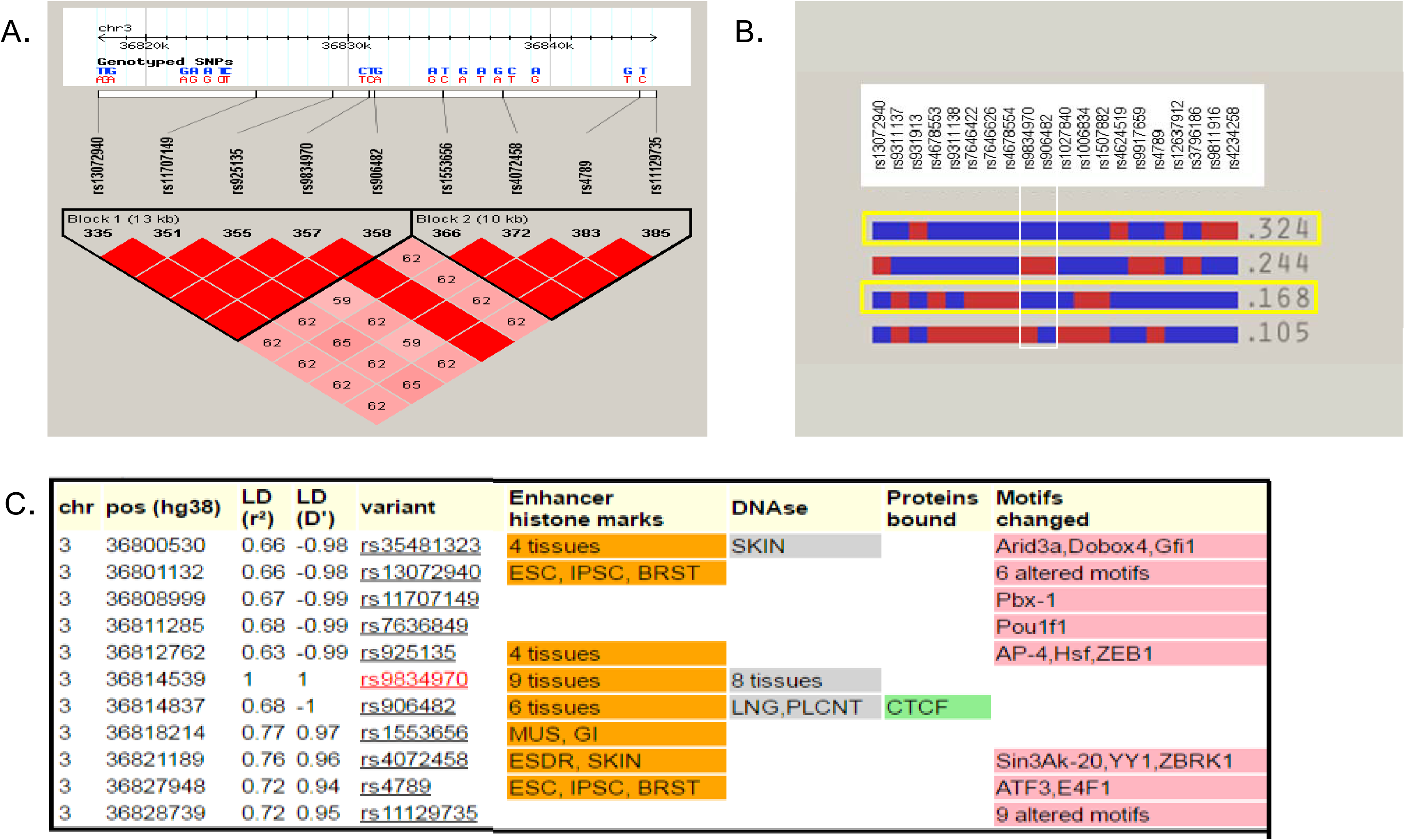
Linkage disequilibrium relationships and functional annotation of common SNPs in GWAS-implicated region upstream of *TRANK1*. (A). Linkage disequilibrium (LD) plot. Common SNPs in the region fall into two large haplotype blocks. LD calculated as D’ in HapMAP Phase III CEU samples using Haploview 4.1; haplotype blocks based on confidence intervals (Zhu et al., 2004). Red squares depict D’ values of 100%; pink squares show D’ values <100%. (B.) Block 1 haplotypes with CEU frequencies >5%. Both common haplotypes that carry the risk allele at rs9834970 (outlined in yellow) also carry the G-allele at of rs906482. (C) Functional annotation of common SNPs in LD with rs9834970. SNPs with r^2^>0.6 in CEU were annotated with the core 15-state (ChromHMM) model using HaploReg v4.1. Several SNPs alter chromatin state, DNAse I sensitive sites, or regulatory motifs. Only rs906482 alters protein binding by CTCF. Abbreviations: ESC, embryonic stem cells; IPSC, induced pluripotent stem cells; BRST, mammary epithelial cells; SKIN, foreskin keratinocyte; LNG, fetal lung; PLCNT, placenta; MUS, fetal muscle leg; GI, small intestine; ESDR H1-derived neuronal progenitor cells.

Functional annotation by the Roadmap Epigenomics and ENCODE projects (Ward and Kellis, 2016) shows that most of the SNPs in this region are predicted to have an impact on chromatin state or transcription factor binding in various tissues (Figure 4C), but only rs906482 has been shown to alter DNA-protein binding. This SNP alters binding by the zinc finger protein, CTCF, in multiple cell types (http://insulatordb.uthsc.edu). Since CTCF is one of the most important gene regulatory proteins in vertebrates (Ghirlando and Felsenfeld, 2016), we decided to further investigate the impact of rs906842 on CTCF binding in NPCs.

Electrophoretic mobility shift assays (EMSA) were performed for each of the 2 alleles at rs906482, using biotin-labeled double-stranded oligonucleotide probes comprising 30 bp of genomic sequence surrounding the variant site (probe sequences shown in Table 5S). The results showed that CTCF peptide directly interacted with this genomic region, displaying significantly higher affinity for the (G) allele at rs906482 (Figure 5). As noted above, this allele usually resides on the same haplotype as the risk (G) allele of rs9834970 (Figure 4B). These results suggest that common genetic variation in this region affects expression of *TRANK1* through a CTCF-mediated mechanism.

**Figure 5.**
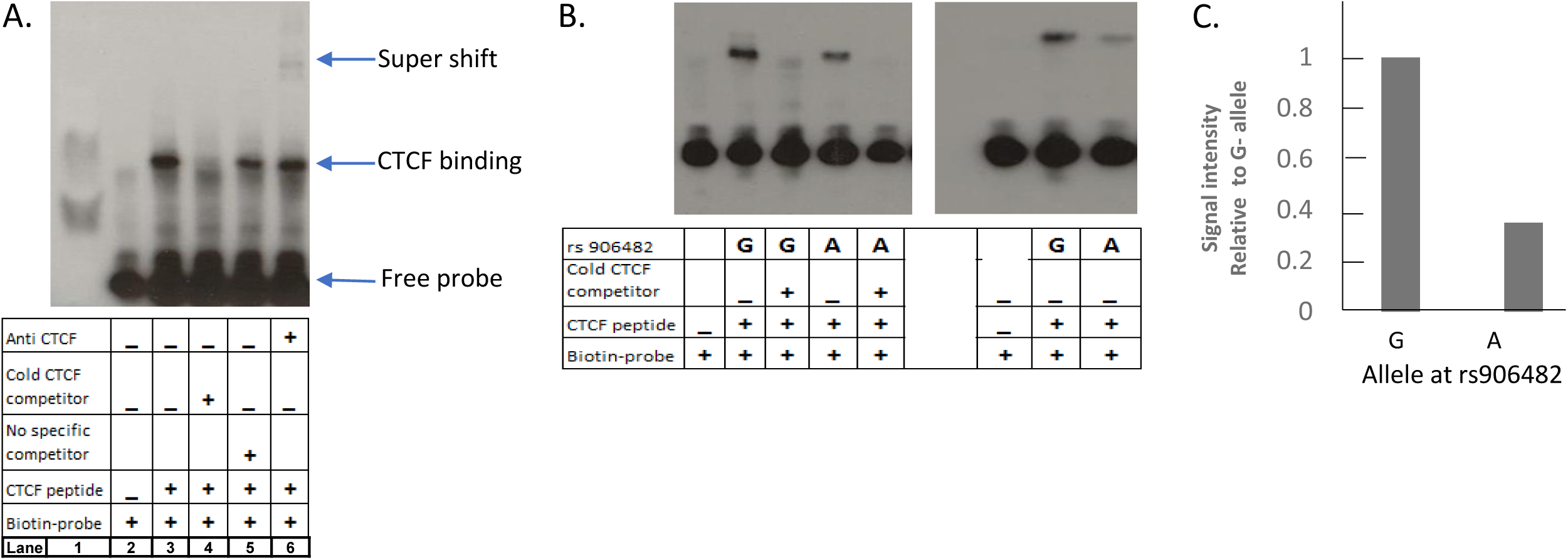
Allele-specific effects on binding of CTCF protein to DNA. (A.) Representative EMSA plot of CTCF binding around rs906482. Lane 1 shows MW marker, lane 2 shows probe-only (negative) control, lane 3 shows probe plus CTCF peptide (positive) control, lane 4 shows that excess unlabeled (“cold”) CTCF competitor displaced binding of biotin-labeled CTCF, lane 5 shows binding of biotin-labeled CTCF with no competitor, lane 6 shows increased MW band due to binding by anti-CTCF antibody (“supershift”). (B.) Genotype-specific differential DNA binding by CTCF. Lane 1 shows probe-only (negative) control, lane 2 shows CTCF binding around G-allele, lane 3 shows displacement by excess unlabeled (“cold”) competitor, lane 4 shows CTCF binding around A-allele, lane 5 shows displacement by unlabeled (“cold”) competitor, lanes 6 to 8 show replicate assays corresponding to lanes 1, 4, and 2. (C.) Quantification of signal intensities from B., lanes 7 and 8. Statistical significance was tested by Student’s t-test.

In light of these results, we repeated the eQTL analyses with genotype at rs906842 as the independent variable. As expected, the results were similar to those observed for rs9834970 (Fig 3), with an even stronger baseline relationship between genotype and *TRANK1* expression. As with rs9834970, VPA also rescued *TRANK1* expression in carriers of the G-allele at rs906842 (Figure S5).

### Genome-wide gene expression profiles

To better understand the impact of decreased *TRANK1* expression at the cellular level, two stable knockdowns of *TRANK1* were generated with *TRANK1* shRNA lentiviral constructs and tested in HeLa cells. The extent of *TRANK1* knockdown was assessed at both mRNA and protein levels, and was found to be substantial (Figure 6A). *TRANK1* mRNA was expressed in the *TRANK1* shRNA knockdown lines at levels 19-24% of those seen in negative control lines transfected with non-target shRNA constructs. *TRANK1* protein was expressed at 21-29% of control levels (Figure 6B).

**Figure 6.**
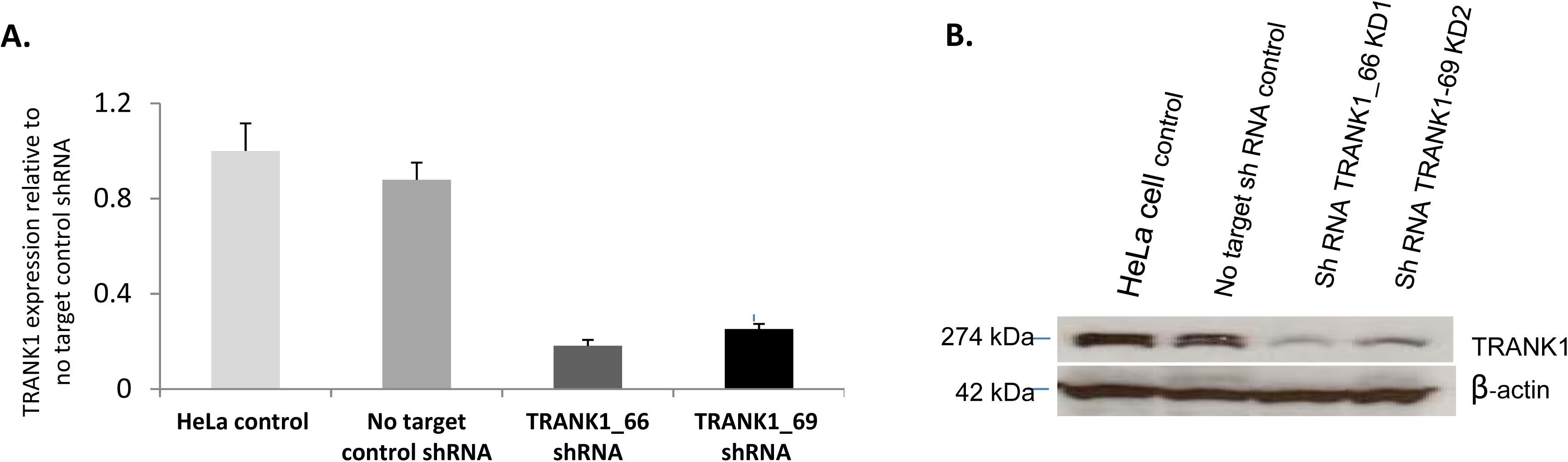
Knockdown of TRANK1 protein expression by shRNA. (A)TRANK1 protein expression in knockdown cell lines, shRNA TRANK1_66 and shRNA TRANK1_69, relative to no-target (scrambled) shRNA. Values based on average of three Western blots. Baseline expression in HeLa lines is shown on the left. (B) Representative Western blot.

Genome-wide gene expression profiling of knockdown cell lines revealed a large number of differentially expressed genes. A total of 214 genes that were differentially expressed with an absolute fold change >1.75 at an FDR<0.05 (Table S1) were enriched for pathways associated with increased cell proliferation and survival and decreased apoptosis (Table 1A). Gene set enrichment analysis indicated that these differentially expressed genes were strongly enriched for several related Gene Ontology (GO) terms (Table S2), including regulation of cell proliferation, multicellular organismal development, and system development.

**Table 1.** Cellular growth and proliferation differences in (A). *TRANK1* shRNA KD in HeLa cell line, (B). VPA treated on iPSC derived neural progenitor cells.

|  |  |  |  |  |  |
| --- | --- | --- | --- | --- | --- |
| A. | Diseases or Functions Annotation | p-Value | Predicted Activation State | Activation z-score | # Molecules |
|  | proliferation of cells | 1.08E-17 | Increased | 2.264 | 93 |
|  | proliferation of lymphocytes | 1.14E-08 | Increased | 2.088 | 27 |
|  | proliferation of neuronal cells | 4.02E-05 | Increased | 2.598 | 18 |
|  | stimulation of cells | 2.04E-12 | Increased | 2.028 | 21 |
|  | Attachment of cells | 3.16E-06 | Increased | 2.121 | 9 |
|  | Cell survival | 1.98E-10 | Increased | 2.140 | 43 |
|  | Apoptosis of muscle cell lines | 4.82E-08 | Decreased | -2.759 | 8 |
| B. | Diseases or Functions Annotation | p-Value | Predicted Activation State | Activation z-score | # Molecules |
|  | formation of cells | 2.76E-04 | Decreased | -2.039 | 29 |
|  | proliferation of Schwann cells | 3.23E-04 |  | -1.948 | 4 |
|  | proliferation of cells | 8.10E-15 |  | -1.793 | 109 |
|  | formation of connective tissue cells | 3.08E-04 |  | -1.622 | 10 |
|  | growth of axons | 3.75E-07 |  | -1.300 | 12 |

Since VPA treatment increases *TRANK1* expression, we expected that VPA treatment of iPSC-derived NPCs would reveal a gene expression profile that contrasted with that seen in the sh*TRANK1* knockdown. A total of 304 genes differentially expressed with an absolute fold change >1.75 at an FDR<0.05 were analyzed (Table S3). As expected, differentially expressed genes in VPA-treated NPCs were enriched for decreased proliferation of cells and decreased growth of axons (Table 1B), in contrast to that seen in the sh*TRANK1* knockdown lines. Similarly, gene set enrichment analysis revealed highly significant enrichment for GO terms related to cell differentiation, nervous system development, neurogenesis, and neuron differentiation (Table S4). Consistent with these results, we observed that treatment of NPCs with VPA substantially decreased cell proliferation (Figure S8.)

### Increasing TRANK1 expression during cellular differentiation and inverse relationship with expression of histone deacetylases

It is notable that among the genes whose expression was increased most by *TRANK1* knockdown (Table S1) was the gene encoding histone deacetylase 1 *(HDAC1)*, which along with other histone deacetylases is known to be inhibited by VPA (Hasan et al., 2013). To more fully characterize the relationship between *TRANK1* and *HDAC1*, we compared relative *HDAC1* and *TRANK1* expression in iPSCs, NPCs, and neurons. *TRANK1* expression was highest in neurons, and lower in iPSCs and NPCs (Figure 7A). In contrast, *HDAC1* and *HDAC2* expression was higher in iPSCs and NPCs than in astrocytes or neurons (Figure 7B-7C). *TRANK1* expression levels rose steadily during neuronal maturation (Figure 7D). In contrast, knockdown of *TRANK1* in HeLa cells led to a substantial increase in *HDAC1* expression (Figure 7E). These data suggest a reciprocal relationship between *TRANK1* expression and *HDAC1* in human cell lines.

**Figure 7.**
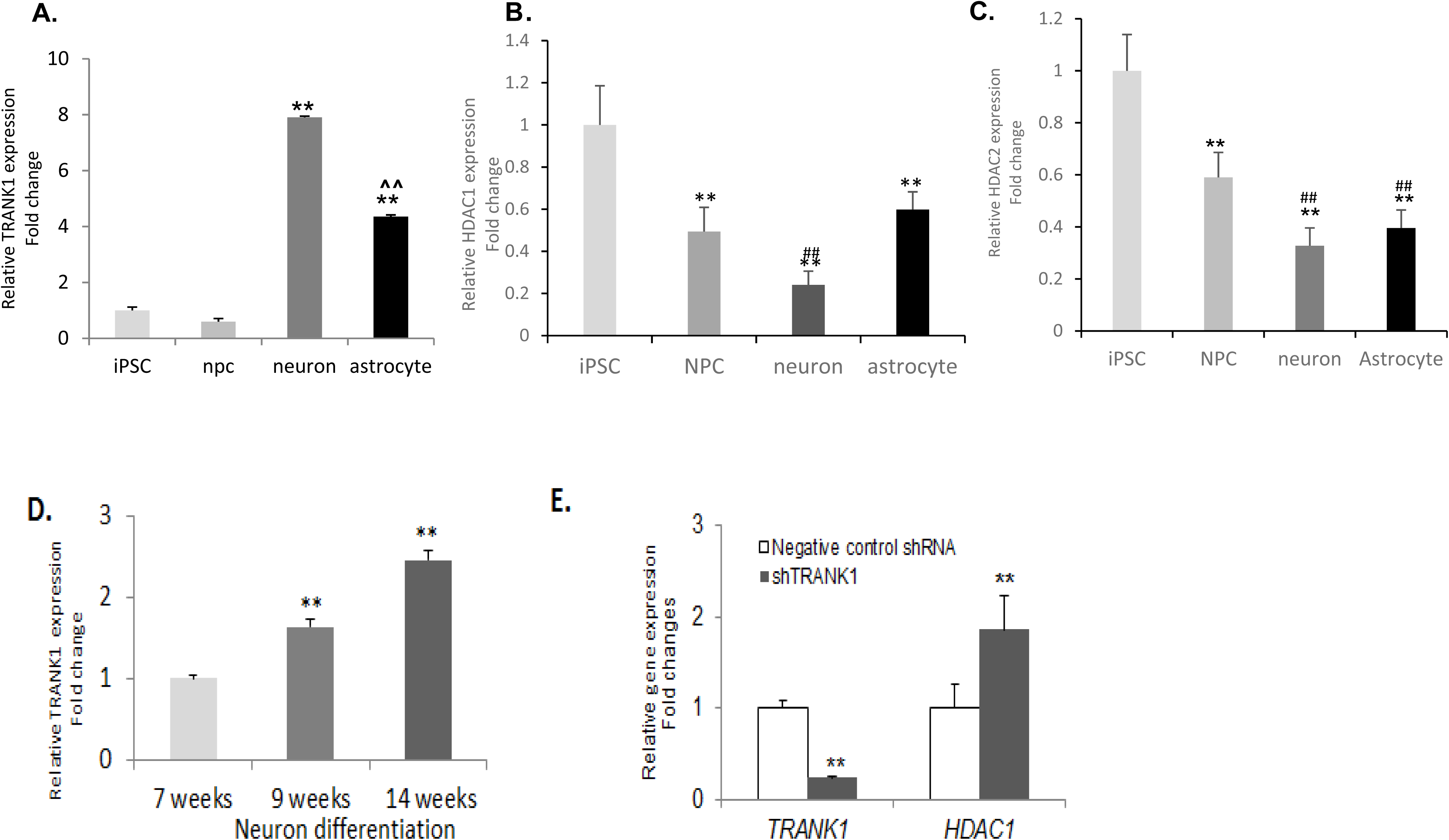
Inverse relationship between expression of *TRANK1* and *HDAC1* or *HDAC2*. Relative mRNA expression of *TRANK1* (A.), *HDAC1* (B.) and *HDAC2* (C.) in iPSC (n=4), NPC (n=11), neurons (n=5), and astrocytes (n=2). **p<0.01, compared to iPSCs; ^^p<0.0.01, compared to neurons; ##p<0.01, compared to NPCs. D. Increase in *TRANK1* mRNA expression during 14 wks of neuronal differentiation and maturation. **p<0.01, compared to 7-wk neurons. E. Relative *HDAC1* and *TRANK1* expression in *TRANK1* knockdown HeLa cells (n=6). **p<0.01, compared to negative control.

## DISCUSSION

In this study, we used human iPSCs and their neural derivatives to investigate the impact of genetic variation and drug treatment on gene expression at a locus implicated by GWAS of major mental illnesses. The results demonstrate a clear link between common SNPs at the locus and expression of mRNA and protein by a nearby gene, *TRANK1*, in neural cells. The results also demonstrate, for the first time, that the bipolar disorder medication, VPA, rescues *TRANK1* expression in neural cells carrying risk alleles at this locus. Although rs9834970 has no known function, a nearby SNP, rs90832, strongly affects binding by the transcription factor, CTCF. Decreased expression of *TRANK1* perturbed expression of many genes involved in neural development and differentiation.

This study has some limitations. The number of iPSC-derived cell lines was limited, reducing power to detect subtle effects. Additional signals may emerge as sample size increases with the inventory of available iPSC. We focused initially on rs9834970, since this SNP had been repeatedly identified as the most associated SNP by previous GWAS of both BD and schizophrenia. However, our findings suggest that a nearby SNP, rs906482 may account for the decreased *TRANK1* expression observed in risk allele carriers. Both SNPs reside upon a haplotype that carries a number of common variants related to chromatin regulation and transcription factor binding (Figure S6), so it is possible that additional SNPs contribute to the biological impact of the risk haplotype. GWAS studies have also identified other SNPs associated with BD and schizophrenia in the *TRANK1* region (2014; Muhleisen et al., 2014; Ruderfer et al., 2013, 2014a; Shinozaki and Potash, 2014). Some of these SNPs are in strong linkage disequilibrium with rs9834970, so may also affect expression of *TRANK1* (Figure S6), but others are not. Further studies that use genome editing techniques such as CRISPR/Cas9 would be needed to establish the functional impact of each individual SNP in this region. It is also possible that variants within the locus affect additional genes that were not fully investigated in the present study.

Drugs can be an important influence on gene expression. This study focused on VPA since it is an effective therapeutic agent for bipolar disorder and demonstrated robust effects on *TRANK1* expression in previous studies (Chen et al 2013a). However, the other effective therapeutic agent we studied, lithium, had no apparent impact on *TRANK1* expression. The *TRANK1* region has also been associated with schizophrenia, and VPA is not known to exert a therapeutic effect in patients with that mental illness. Further studies will be needed to explore the full range of drugs that affect expression of *TRANK1* in neural cells.

These results raise the possibility that the well-established therapeutic effects of VPA in BD may be due in part to its ability to regulate expression of *TRANK1*. VPA rescued the decrease in *TRANK1* expression in cells carrying the risk haplotype. Moreover, *TRANK1* regulated many of the same cellular growth and differentiation pathways that are known to be affected by VPA, highlighting a key role for histone deacetylases, known targets of VPA. VPA affected *TRANK1* expression mainly in less mature cells, consistent with the observation that exposure to VPA can be teratogenic during early stages of nervous system development (Kim et al., 2011; Yu et al., 2009). However, it is not obvious from these data how VPA would exert a beneficial effect in the mature brain. VPA did have a significant impact on *TRANK1* expression in astrocytes, suggesting one way that the therapeutic window could extend into adulthood. Recent work has highlighted the potential importance of astrocytes and other glial cells in the pathophysiology of a variety of neuropsychiatric disorders.(Chung et al., 2015; Peng et al., 2016; Stevens and Muthukumar, 2016)

This study demonstrated reciprocal effects of *TRANK1* and *HDAC1* on gene expression, cell proliferation, and cell differentiation. We observed a significant increase of *TRANK1* expression over baseline in more differentiated cells, where *HDAC1* expression was decreased. Future studies might assess whether similar cellular phenotypes follow treatment with other *HDAC* inhibitors. Our results are consistent with the emerging literature implicating VPA and other histone deacetylase inhibitors in neuronal growth and differentiation (Ganai et al., 2016; Gottlicher et al., 2001; Yu et al., 2009).

While the precise mechanism whereby VPA and noncoding genetic variation near *TRANK1* affect gene expression remains to be fully elucidated, our findings suggest an important role for the multi Zn-finger protein, CTCF. We have shown that genetic variation within risk haplotypes dramatically alters CTCF binding affinity, and that decreased CTCF binding at this locus correlates with increased expressed of *TRANK1*. CTCF influences gene expression by acting as a transcriptional insulator, orchestrating long-range DNA-looping interactions between distal enhancers and their cognate promoters to activate or repress gene expression. (Guo et al., 2015). Genome-wide analysis of CTCF has demonstrated its crucial role in higher-order chromatin organization that can increase or curtail enhancer-promoter interactions, depending on the relative positions of these regulatory elements (Ghirlando and Felsenfeld, 2016). Increased CTCF binding to the risk allele may block the effect of a distal enhancer that otherwise activates *TRANK1* expression. Finding that enhancer is an important future goal and may suggest other therapeutic approaches to the regulation of *TRANK1* expression in the brain.

We speculate that CTCF may also play a role in the mechanism whereby VPA impacts *TRANK1* expression. Previous studies have shown that CTCF can recruit histone deacetylases to the transcription activation complex (Lutz et al., 2000) and that VPA is a potent histone deacetylase inhibitor (Gurvich et al., 2004). VPA also decreases CTCF expression (Lutz et al., 2000; Oti et al., 2016). Thus, VPA may partially antagonize gene repression by CTCF at the *TRANK1* locus.

The approach we have taken illustrates a potential strategy to investigate GWAS findings in neuropsychiatric disorders. Genotype-specific iPSC-derived cells can be used to study the relationships between allelic variation and gene expression (or splicing) across a range of cell types, developmental stages, and environmental conditions, offering a degree of cellular, temporal, and experimental resolution that is not possible with blood or post-mortem brain studies. Recent studies have demonstrated that the effects of cis-acting regulatory variation can differ between brain regions (Buonocore et al., 2010), developmental stages (Burkhardt et al., 2013), and cell types (Won et al., 2016). Human iPSCs and their neural derivatives also provide a valuable experimental platform for screening therapeutic drugs in patient-specific cells. We have shown how such studies can identify potentially causal genetic variants, enhance understanding of disease mechanisms, and illuminate relationships between genetic variation and treatments for neuropsychiatric disorders. More work is needed to develop scalable, high-throughput strategies for screening larger numbers of genetic risk variants, especially variants whose regulatory impact is specific to mature cells, such as neurons, or variants that affect genes involved in multi-cellular functions, such as neural circuits.

## EXPERIMENTAL PROCEDURES

### Generation and Reprogramming of iPSCs

Eleven iPSC lines were used in this study (Table 2). Seven human iPSC lines were obtained from the NIH (Bethesda, Maryland), five of which were reprogrammed at the NHLBI Stem Cell Core and two of which were reprogrammed at the NIH Center for Regenerative Medicine (CRM). Lines GM05990 and 10593 were reprogrammed in our laboratory (HGB) from fibroblasts obtained from Coriell Cell Repositories or from a consented participant in our ongoing study of bipolar disorder (10593; obtainable from the Rutgers University Cell and DNA Repository as 10C117904). Lines GM23240 and GM23476 were obtained as iPSC from Coriell Cell Repositories. Most iPSCs in this study were generated using lentiviral transduction cells with STEMCCA vector (Millipore, Billerica, MA).

**Table 2.**
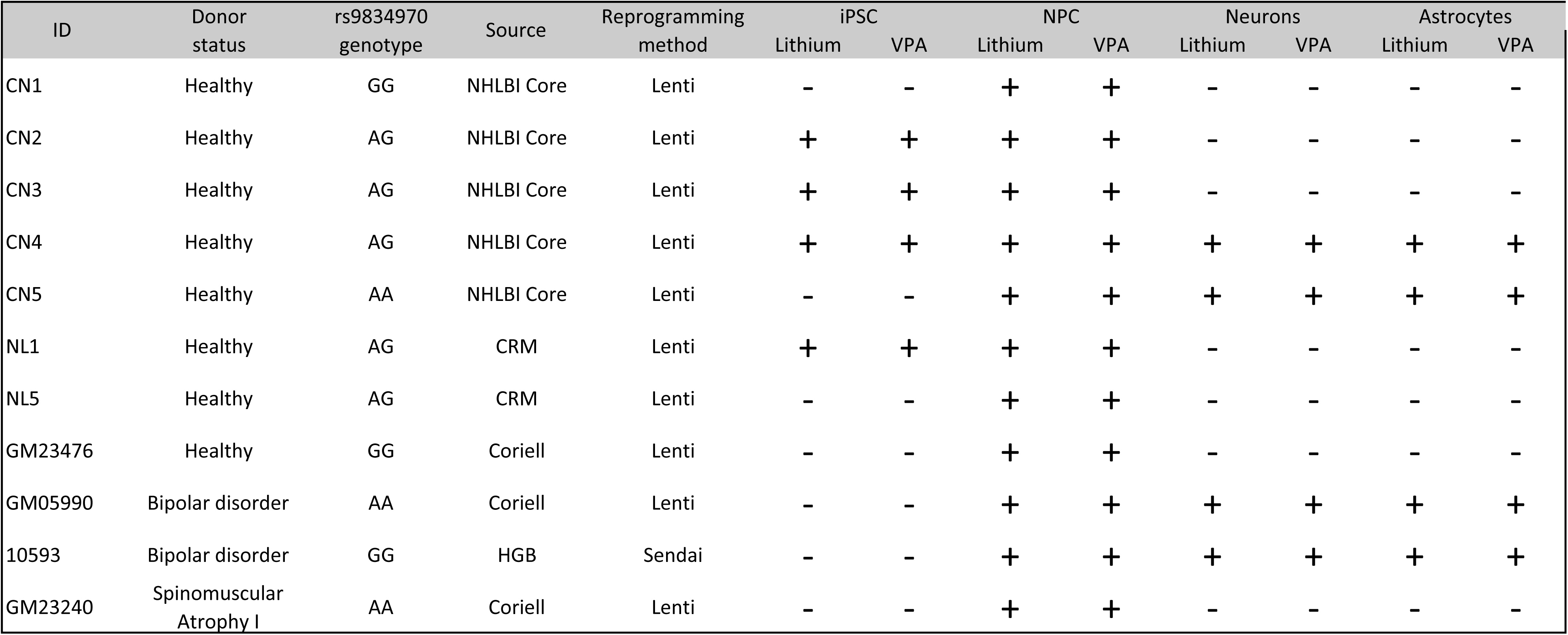
Cell lines used in present study

| ID | Donor status | rs9834970 genotype | Source | Reprogramming method | iPSC |  | NPC |  | Neurons |  | Astrocytes |  |
| --- | --- | --- | --- | --- | --- | --- | --- | --- | --- | --- | --- | --- |
|  |  |  |  |  | Lithium | VPA | Lithium | VPA | Lithium | VPA | Lithium | VPA |
| CN1 | Healthy | GG | NHLBI Core | Lenti | - | - | + | + | - | - | - | - |
| CN2 | Healthy | AG | NHLBI Core | Lenti | + | + | + | + | - | - | - | - |
| CN3 | Healthy | AG | NHLBI Core | Lenti | + | + | + | + | - | - | - | - |
| CN4 | Healthy | AG | NHLBI Core | Lenti | + | + | + | + | + | + | + | + |
| CN5 | Healthy | AA | NHLBI Core | Lenti | - | - | + | + | + | + | + | + |
| NL1 | Healthy | AG | CRM | Lenti | + | + | + | + | - | - | - | - |
| NL5 | Healthy | AG | CRM | Lenti | - | - | + | + | - | - | - | - |
| GM23476 | Healthy | GG | Coriell | Lenti | - | - | + | + | - | - | - | - |
| GM05990 | Bipolar disorder | AA | Coriell | Lenti | - | - | + | + | + | + | + | + |
| 10593 | Bipolar disorder | GG | HGB | Sendai | - | - | + | + | + | + | + | + |
| GM23240 | Spinomuscular Atrophy I | AA | Coriell | Lenti | - | - | + | + | - | - | - | - |

The process for iPSC generation included reprogramming human fibroblasts into iPSCs using the four classic Yamanaka transcription factors (OCT4, SOX2, KLF4, and c-MYC) (Kulzer et al., 2014; Takahashi et al., 2007). Transduced human fibroblasts were initially grown over X-irradiated mouse embryonic fibroblasts (MEFs) (MTI-Global Stem(Gaithersburg, MD) on 1% gelatin coated plates in the human embryonic stem cell (hESC) medium (Dulbecco’s modified Eagle’s medium/F-12 containing 20% KnockOut serum replacer, 0.1 mM non-essential amino acids, 1 mM L-glutamine (all from Life Technologies, Grand Island, NY), 0.1 mM β-mercaptoethanol (Sigma-Aldrich, St. Louis, MO), supplemented with 4 ng/ml bFGF (Millipore, Billerica, MA). Colonies with human embryonic stem cell (hESC)-like morphology emerged within 10 to 14 days after fibroblast reprogramming, and displayed the typical morphology of iPSCs by 20 days. Three to four weeks after reprogramming, colonies were manually picked and individually passaged onto irradiated MEF feeder cells. After a few passages in MEF feeder cells with StemPro medium (Invitrogen, Grand island, NY), iPSC colonies were switched to 1% Matrigel-coated plates (BD Biosciences, Franklin Lakes, NJ) with E8 medium (Life Technologies, Grand Island, NY) and then switched to mTeSR1 (StemCell Technologies, Vancouver, Canada), Cells were frozen in freezing medium (mTeSR1, 10% DMSO).

### Monolayer culture of iPSC

In order to reduce variation in cell number and better equalize exposure to drug treatments, four colonies of iPSCs were adapted to single-cell based non-colony type monolayer (NCM) culture as described (Chen et al., 2014; Kulzer et al., 2014). Briefly, iPSC colonies were dissociated with Accutase (Life Technologies, Grand Island, NY), approximately 1.3 to 2×10^6^ hESCs were seeded in 6-well plates coated with 2% Matrigel in E8 medium with 10 μM ROCK inhibitor (TOCRIS Bioscience, London, UK). After three hours, the medium was replaced with E8 medium without ROCK inhibitor. The cells were allowed to grow as a single cell monolayer for three to four days, and the medium was changed daily. Treatment was given 24 hours after cell plating.

### Generation of NPCs and differentiation into neurons and astrocytes

Eleven neural precursor lines were used in this study. Two lines (NL1 and NL5) were obtained from the NHLBI Stem Cell Core. Nine iPSC lines were differentiated into NPCs in-house using a published neural precursor cell generation method (Kozhich et al., 2013) and Gibco PSC neural induction medium (Life Technology, Carlsbad, CA). Briefly, NPCs were generated by first differentiating iPSCs to embryoid bodies from AggreWell™ 800 system (StemCell Technologies, Vancouver, Canada) in STEMdiff™ neural induction medium; this allows 800 EBs of uniform size to form in a week. After 10-12 days, neural tube-like rosettes were detached mechanically, the embryoid bodies in suspension were collected and plated on polyornithine/laminin-coated plates in STEMdiff™ Neural Progenitor Medium (StemCell Technologies, Vancouver, Canada). Neuronal differentiation was induced by changing the neural progenitor medium to neuron culture medium consisting of Neurobasal (Invitrogen, Grand island, NY), supplemented with 1× B27 (Invitrogen), 10 ng/ml brain derived neurotrophic factor (BDNF) (Peprotech, Rocky Hill, NJ), and 10 ng/ml glial cell line-derived neurotrophic factor (GDNF) (Peprotech, Rocky Hill, NJ), L-ascorbic acid (200 ng/ml), cAMP (1μM) (Sigma-Aldrich, St. Louis, MO). The neuron medium was refreshed every three to four days. The cells were cultured in neuron medium for more than two months. To differentiate NPCs into astrocytes, NPCs were plated on a Geltrex-coated culture dish in STEMdiff™ Neural Progenitor Medium. After 2 days, medium was changed to DMEM, 1% N2 supplement (Invitrogen, Grand island, NY), 1% FBS, 20 ng/ml CTNF (R&D Systems, Minneapolis, MN) medium. In 6 to 8 weeks, 60-80% cells were positive for astrocyte-specific markers.

### Human iPSC characterization

Human iPSCs were rigorously characterized using several assays. Immunocytochemistry was performed for different pluripotency markers. Pluripotency was also assessed by immunostaining for surface and nuclear pluripotency markers, flow cytometry quantification, quantitative reverse transcription (RT)-PCR of endogenous pluripotency genes, and by gene-chip and bioinformatics-based PluriTest assays (Illumina, San Diego, CA). The properties of iPSC-derived NPCs, neurons, and astrocytes were characterized by standard immunochemical analysis, qRT-PCR, electrophysiology, and gene-chip and bioinformatics-based PluriTest assays. iPS clones that displayed a normal karyotype by either G-banding or spectral karyotyping (NHGRI Cytogenetics Core Facility) were selected for further studies.

### Immunocytochemistry

Immunocytochemistry was performed as previously described (Kozhich et al., 2013), iPSCs and neural derivatives were fixed in 4% paraformaldehyde for 15 min and washing 3 times in PBS. Pluripotency of iPSC were detected using mouse anti-Oct-3/4, anti-SSEA1, rabbit anti-NANOG), mouse anti-SSEA-4 (all from Sigma-Aldrich, St. Louis, MO), mouse anti-TRA1-60 or mouse anti-TRA1-81 (Santa Cruz Biotechnology, Dallas, Texas). Cell nuclei were counterstained with DAPI.

Primary antibodies for immunostaining of NPCs included SOX2 (1:500, R&D Systems, Minneapolis, MN),), Nestin (1:500, Millipore, Billerica, MA), and Pax6 (1:200, Abcam, Cambridge, MA). Mouse anti-Beta-tubulin-III (TUJ1) (Millipore, Billerica, MA), rabbit anti-MAP2 (Millipore, Billerica, MA), rabbit anti-glial marker GFAP (StemCell Technologies, Vancouver, BC, Canada), mouse anti-A2B5 (1:1000; Sigma-Aldrich, St. Louis, MO), mouse anti-tyrosine hydroxylase (TH) (StemCell Technologies, Vancouver, BC, Canada), and rabbit anti-GAD65/67 (Sigma-Aldrich, St. Louis, MO). These antibodies were used to identify differentiated neurons, astrocytes, oligodendrocytes, glutamatergic neurons, and GABAergic neurons. The secondary antibodies were Alexa488 anti-mouse, Alexa 555 anti-mouse, Alexa488 anti-rabbit, Alexa555 anti-rabbit (at 1:1000 concentration; Life Technologies, Grand Island, NY). Images were acquired by use of a fluorescence microscope (EVOS). Rabbit anti-postsynaptic density 95 (PSD95) (1:1000, Abcam, Cambridge, MA), a postsynaptic density marker, was used to stain differentiated neurons, and images were collected with a confocal microscope (Zeiss, Oberkochen Germany). Rabbit anti *TRANK1* (1:400, Sigma-Aldrich, St. Louis, MO) was used for Western blot analysis.

### Electrophysiology

For electrophysiological recordings, cells were grown on 18 mm glass coverslips coated with polyornithine/Laminin in 24-well plates. Cells were investigated six to 12 weeks after neuron differentiation began. The coverslips were transferred to a submerged chamber. Single cell with a bright cell body, and two or more neurites was identified morphologically under an Olympus BX51WI microscope equipped with infrared differential interference. Recordings were performed at room temperature under continuous perfusion (1.5 ml / min) with artificial cerebrospinal fluid (ACSF) bubbled with 5% CO2 and 95% O2, containing (in mM) NaCl 119, KCl 2.5, NaHCO3 26.2, NaH2PO4 1, glucose 11, CaCl2 2.5, and MgSO4 1.3. The osmolarity of the ACSF was adjusted to 300 mOsm and pH was adjusted to 7.4. Patch pipettes (4 – 6 MΩ) were filled with intracellular solution containing (in mM): 130 KMeSO4, 10 KCl, 10 HEPES, 4 NaCl, 4 Mg-ATP, 0.3 NaGTP and 1 EGTA with osmolarity and pH at 285 mOsm, 7.3 respectively. Data for action potential were collected under current-clamp mode a Multiclamp 700B amplifier (Axon Instruments, Foster City, CA), filtered at 2 kHz, and digitized at 50 kHz. Resting membrane potentials were determined immediately after gaining whole-cell access. Action potentials were elicited by applying increasing depolarizing current pulses (10 pA current steps). For voltage-clamp recordings, cells were clamped at −70 mV; Na^+^ currents and K^+^ currents were evoked by voltage step depolarization. Command voltages varied from −80 to + 10 mV in 10 mV increments.

### DNA and RNA Extraction and Quantitative real time PCR (qPCR)

Genomic DNA was prepared using the DNA Mini Kit (QIAGEN, Valencia, CA). Total RNAs were extracted using the RNeasy kit (QIAGEN, Valencia, CA) per the manufacturer’s instructions. Genomic DNA contamination was removed using a DNA-free kit (Invitrogen, Carlsbad, CA). Superscript III reverse transcriptase (Invitrogen, Grand Island, NY) was used for cDNA synthesis. mRNA levels of selected genes were determined using Roche LightCycler 480 and Roche Universal ProbeLibrary System (Roche, Basel, Switzerland). The comparative *C*T method (2^−δ*C*^T method) was used to quantify relative mRNA levels. *TRANK1* gene expression levels were measured by q-PCR, with three technical replicates for each treatment condition. Sequence information for all primers used for qPCR is in Supplementary Table 5.

### Cultured hippocampal neurons

Hippocampal neurons were prepared from fetal rat brain at embryonic day 18 (E18). Dissected pieces of hippocampi were incubated with 0.5% trypsin, followed by 2mg/ml trypsin inhibitor (Sigma, St. Louis, MO), then dissociated mechanically. Following centrifugation, cells were re-suspended in Neurobasal medium, supplemented with B27 (1x) and glutamine (2 mM), and cultured on 6 well plates coated with poly-D-lysine (Sigma, St. Louis, MO). Three-quarters of the culture medium was replaced every 4 days. Drug treatment was performed after 8 days of culture.

### Drug Treatment

Valproic Acid (VPA) (Sigma, St. Louis, MO) was diluted in sterile PBS to prepare a 500 mM stock solution and frozen in 220 μl for up to two weeks. Lithium chloride (Sigma, St. Louis, MO) was diluted in sterile PBS to prepare a 1M stock solution. Cells were treated with lithium chloride (1 mM), VPA (0.5 mM), or VPA (1mM). These concentrations are within typical therapeutic ranges for treatment of bipolar disorder. Treatments were applied for 72 hours in four iPSC, eleven NPC, five neuronal, and two astrocyte cell lines, each with 3 technical replicates.

### Western Blotting

Protein isolation was performed with M-PER lysis buffer (Thermo scientific, Catalog 78503). Concentration was determined using a spectrophotometer. Twenty-five micrograms of protein were loaded onto 4–12% Tris-Glycine gels, electrophoresed, and transferred with iBlot 2 onto nitrocellulose membranes (Life Technologies, Grand Island, NY). The membrane was blocked, incubated with anti-TRANK1 antibody washed 5 times with TBST, and incubated with secondary antibody After washing, the membrane was exposed to Amersham™ ECL™ Western blotting detection reagents (GE Healthcare, Pittsburgh, PA) and developed. ImageJ was used for quantifying Western blotting data.

### Karyotypes

iPSC clones were prepared for Spectral karyotyping (SKY) following the method described by Moralli et al. 2011. {Moralli, 2011 #23822}Stem Cell Rev and Rep (2011) 7:471-477). SKY was performed to visualize all human chromosomes, each individually colored with a different fluorescent probe (Applied Spectral Imaging Inc., Carlsbad, CA).

### Fluorescent activated cell sorting (FACS) analysis

Each of two neural progenitor cell lines were seeded into a 6-well plate at a density of 1x10^5^ or 1x10 ^4^ cells per well and incubated overnight prior. Cells were treated with vehicle or 0.5mM VPA for 24 hours or eight days, then collected for cell cycle analysis. Flow cytometric cell cycle analysis was performed as previously described (Reddy et al., 1997). Briefly, both treated and untreated cells were washed with PBS, then dissociated with Accutase for 30 minutes at 37°C. Dissociated cells were washed three times in PBS in 2% bovine bovine serum albumin (BSA), counted, re-suspended in 1 ml PBS, and fixed with 70% ethanol overnight at -20°C. Cells were washed with PBS and stained for 40 minutes at 37°C with 50 μg/ml propidium iodide in PBS (containing 200 μg/ml RNase A and 0.1% Triton X-100). Cells were analyzed using a Quanta SC flow cytometer (Beckman Coulter, Indianapolis, IN). After gating out doublets and debris, cell cycle distribution was analyzed using Summit 4.3 (Dako Colorado, Inc., Fort Collins, CO).

### Cell Proliferation and Cytotoxicity Assay

The effect of VPA on cell growth and cytotoxicity were determined by using trypan blue staining to count cell number and viability with an automatic cell counter. 3-(4,5-dimethylthiazol-2-yl)- 2,5-diphenyltetrazolium bromide (MTT; Sigma-Aldrich, St. Louis, MO) absorbance in living cells was also measured, as previously described(Hecker et al., 2010). In brief, equal amounts of NPCs (5x10^5^ cells/well in 6 well plates for cell counting and 3000 cells/well in 96-well plates (Nunc) for MTT), were seeded and incubated overnight before VPA treatment. After exposure to the designated doses of VPA for the indicated times, MTT solution [20 μl: 2 mg/ml in phosphate-buffered saline (PBS)] was added to each well of the 96-well plates. The plates were incubated for four more hours at 37°C. Medium was withdrawn from the plates by pipetting and an additional 200 μl DMSO was added to solubilize the crystals. The optical density was measured at 570 nm using a microplate reader (Synergy™ 2, BioTek Instruments Inc, Winooski, VT). Trypan blue staining and automated cell counting (Bio-Rad TC20 automated cell counter, Lenexa, KS) were used to count cells and percentage of live cells in each treatment condition.

### RNA interference

MISSION™ shRNA Lentiviral Transduction Particles (Sigma Aldrich, St. Louis, MO) were used for the knockdown study. First, a puromycin kill curve was performed with each cell line to determine the optimal amount of puromycin needed for selection without toxicity to transduced cells. In HeLa cells, the optimal concentration of puromycin (Calbiochem, La Jolla, CA) was 3 μg/mL. Four gene-specific shRNA sequences were designed for human *TRANK1* gene; one negative construct and one “non- target” construct were transduced separately into HeLa cells (Table S6). The negative construct contained only vector. The “non-target” construct contained a short hairpin RNA sequence that did not code for any known human gene, and acted as a control. HeLa cells were seeded in 48-well plates (50% confluent) overnight, and infected with lentivirus at multiplicity of infection (MOL) of 10, with serum-free, antibiotic-free DMEM media containing 8 μg/mL of Polybrene (Sigma Aldrich, St. Louis, MO). After six hours, the lentiviral shRNA medium was removed and replaced with DMEM-containing 20% FBS. Forty-eight hour post-transfection, the cells were seeded into a 6-well plate and propagated in complete DMEM medium with 20% FBS and 3 μg/ml puromycin until colonies were visible. The puromycin-resistant clones were picked and expanded, and the knockdown efficiency was verified by quantitative RT-PCR and western blot. The two constructs with the best knockdown efficiency (over 70%), along with non-target shRNA control, were used for gene expression array studies.

### Electrophoretic mobility shift assay

Electrophoretic mobility shift assay (EMSA) was performed by using the Gel shift chemiluminescent EMSA kit purchased from Active Motif. Sequences for synthetic double-stranded consensus CTCF and 5’ biotin-labeled oligonucleotides corresponding to the rs906482 are listed in Table S7. The sequence of consensus CTCF double-stranded oligonucleotide was derived from a binding site previously described to bind to CTCF (Xiao et al., 2011). CTCF-associated peptide was affinity purified using the procedure described previously (Yusufzai and Felsenfeld, 2004). Oligonucleotide probes were designed with both alleles of rs906482 flanked by14 bp in both a cold and 5’-biotinylated form (IDT). Biotinylated and un-biotinylated oligonucleotides were annealed with the reverse complementary oligonucleotide (95 °C to 25 °C temperature stepdown). For each binding reaction, 5–10 pmol purified CTCF peptide was incubated for 20 min at room temperature with 20 fmol biotin-labeled duplex oligonucleotide containing either A or G allele in rs906482 with or without consensus CTCF oligonucleotide in binding buffer. After incubation, the mixture was loaded on a 6% DNA retardation gel (EC6365BOX from Life Technologies) and separated by electrophoresis in 0.5 TBE buffer at room temperature (100 V for 1 h and 20 min). Following electrophoresis, samples were transferred to a nylon membrane (380 mA, 30 min). Transferred DNA was then cross-linked using UV-light to membrane and detected by chemiluminescence. Products were detected by stabilized streptavidin–horseradish peroxidase conjugate (Pierce). For competition assays, unlabeled consensus oligonucleotides at 100-fold molar excess were added to the reaction mixture before addition of the biotin-labeled probe. Detected bands were measured and analyzed by computer quantification using Image J Software (https://imagej.nih.gov/ij/).

### Microarray Analysis

Gene expression profiling was performed on RNA extracted from the following samples: a) iPSC-derived NPC lines treated for 72 hr with 0.5mM VPA (n=8) or vehicle (n=8); and b) two HeLa lines each treated with a different shTRANK1 probe for 72-hr. Groups were equally divided between array plates and hybridization batches.

Total RNA was extracted using the RNeasy Mini kit (QIAGEN, Hilden, Germany). Total RNA was used for cRNA amplification, Cy3 labeling, and hybridization on to Illumina HT-12_V4 beadchips (48, 000 probes, Illumina, San Diego, CA) following the manufacturer’s protocol (Illumina, San Diego, CA). Microarray data were assembled using Genome Studio V3.0 (Illumina, San Diego, CA). Data were further analyzed with both Gene Spring software and a Raw expression data were log2 transformed, and quantile normalized using a custom R script calling Bioconductor (www.bioconductor.org). Outliers were checked based on inter-sample correlations and principal component analysis. Transcripts were considered robustly expressed and included in the analysis if the coefficient of variation lay within the linear phase of the distribution (Table S1). Based on this, 12,568 probes passed with VPA treatment as the independent variable. Differential expression was tested by one-way analysis of variance. Differentially expressed genes were identified based on a p<0.05 and an absolute fold-change between conditions of 1.2 (Table S2). Gene-set enrichment analysis was carried out with DAVID (Huang da et al., 2009a, b), at medium stringency, with background specified as the total set of robustly-expressed probes.

### Genotyping and sequence

All samples were genotyped on the Illumina Infinium Human OmniExpress Exome bead array (Illumina, San Diego, CA). Linkage disequilibrium with rs9834870 was evaluated using the tagger software implemented in Haploview version 4.1 (Broad Institute of MIT and Harvard, Boston, Massachusetts).

To further validate genotypes in the NPCs and detect rare variants not represented on the SNP array, 400 bp flanking rs9834970 was Sanger-sequenced at Macrogen. All samples showed the expected genotypes at rs9834970. No additional variants were detected (Figure S7).

### Conditional analysis of genetic association signals in the TRANK1 locus

We used the approximate conditional analysis method implemented in GCTA to conduct association analysis of all markers in the TRANK1 locus conditional on the index SNP rs9834970. Unlike traditional conditional analysis methods that require raw genotype data, this approach uses meta-analysis summary statistics and an LD matrix derived from a reference sample to perform approximate conditional association analysis. In our analysis, we used 3,158 individuals of European ancestry as the reference sample.

## SUPPLEMENTAL INFORMATION

Supplemental Information includes Supplemental Experimental Procedures, eight figures, and seven tables.

## AUTHOR CONTRIBUTIONS

X.J. and F.J.M. conceived the project, designed the experiments, conducted data analyses, and wrote the manuscript. X.J. performed most of the experimental procedures. S.D-W generated the GM05990 iPSC line, N.A. carried out SNP genotyping, X.G. performed and analyzed the electrophysiological experiments, T. X. and G. F. helped to design CTCF and EMSA assays and provided reagents and technical assistance; B.M. provided technical assistance and advice on culture and differentiation of iPSC lines; L.H. performed the conditional association analysis. All co-authors reviewed the manuscript before submission.

## ACKNOWLEDGMENTS

**S**upported by the Intramural Research Programs of the National Institute of Mental Health (NIMH; ZIA-MH00284311/NCT00001174), National Institute of Neurological Disease and Stroke (NINDS), and the National Institute of Diabetes and Digestive and Kidney Diseases, National Institutes of Health. The authors have no conflicts of interest to disclose, financial or otherwise. Dr. Kevin Chen, at the NINDS Stem Cell Unit helped with Flow Cytometry Analysis; Dr. Wei Lu at the Synapse and Neural Circuit Unit (NINDS) helped with electrophysiological recordings; Dr. Mahendra Rao, formerly of the Center for Regenerative medicine (CRM), provided 2 neural progenitor lines; Dr. Manfred Boehm (Laboratory of Cardiovascular Regenerative Medicine, NHLBI) provided 5 iPSC lines; Dr. Kory Johnson (NINDS) Microarray Core helped analyze the microarray gene expression data; Drs. Amalia Dutra and Evgenia Pak of the NHGRI Cytogenetics Core performed spectral karyotyping. GM05990, GM23240, and GM23476 were obtained from Coriell Cell Repositories (Camden, NJ). Line 10593 was obtained from the Rutgers University Cell and DNA Repository (Piscataway, NJ; catalog #10C117904). Special thanks to Ioline Henter (NIMH), who provided invaluable editorial assistance.

